# The proteasome-related Connectase is a fast and specific protein ligase

**DOI:** 10.1101/2020.08.14.217505

**Authors:** Adrian C. D. Fuchs, Moritz Ammelburg, Jörg Martin, Marcus D. Hartmann, Andrei N. Lupas

**Affiliations:** Max Planck Institute for Developmental Biology, Department of Protein Evolution, Max-Planck-Ring 5, 72076 Tübingen, Germany

## Abstract

Genetic methods allow the recombinant production of any protein of interest, but yield the full-length construct in one step and are limited to native amino acids. For the “on demand” generation of chimeric, immobilized, fluorophore-conjugated or segmentally labeled proteins, these proteins must be modified using chemical^1,2^, (split) intein^2,3^, split domain^4^ or enzymatic methods^5^. While each of these options comes with its own advantages and drawbacks, ligase enzymes are often used where small ligation motifs and good chemoselectivity are required. However, applications with the reference enzyme Sortase A are impeded by poor catalytic efficiencies, low substrate specificities and side reactions^6,7^. Here, we present the discovery of Connectase, a monomeric proteasome homolog that ligates substrates via a highly conserved KDPGA motif originally identified in methyltransferase A (MtrA), a key enzyme in archaeal methanogenesis. Connectase displays nanomolar affinity and thus great specificity for its substrates, allowing efficient protein-protein ligations even in complex solutions and at low substrate concentrations. Compared to an optimized Sortase variant, Connectase catalyzes such ligations at substantially higher rates, with higher yields but without detectable side reactions and thus presents a valuable new tool for protein conjugations.

## Connectase binds and modifies MtrA

The proteasome is a large multi-subunit barrel-shaped complex that plays an important role in eukaryotic cells as the main protease in the targeted protein degradation pathway. While studying the function of new, yet uncharacterized proteasome homologs, we identified a representative denoted as Domain of Unknown Function (DUF) 2121 in public databases and as Connectase in the following. Connectase shares only about 11% sequence identity with proteasomal subunits but assumes a similar fold in structure predictions and retains the putative active site Ser/Thr residue in position 1 (Extended Data, Fig. 1). Representatives are found in most archaea capable of producing methane from carbon dioxide and molecular hydrogen (hydrogenotrophic methanogenesis^8^; Extended Data, Fig. 2), with optimal growth temperatures ranging from 25°C – 110°C^9^. Amongst those, two Connectase variants exist: in class II methanogens^10^, such as *Methanosarcina mazei*, the protein is composed of just the regular proteasome-type NTN-domain (N-terminal nucleophile), whereas in class I methanogens, such as *Methanocaldococcus jannaschii*, it features an extra small C-terminal beta-barrel domain with remote similarity to OB-folds (oligosaccharide binding). Although the speciation event separating class I and II methanogens has occurred probably more than 3 billion years ago^11^, both mesophilic *M. mazei* and thermophilic *M. jannaschii* variants (36% sequence identity) showed similar characteristics in our experiments and both could be produced as soluble proteins (> 20 mg/ml) in *E. coli* with high yield (> 40 mg per liter of culture).

Mass spectrometric analysis of a pulldown experiment with *M. mazei* Connectase using whole cell extract resulted in the identification of a potential interactor, methyltransferase A, which is part of the membrane-bound MtrA-MtrH complex and acts in the hydrogenotrophic methanogenesis pathway (Extended Data, Fig. 3a;^12^). This interaction was confirmed by co-elution of a purified MtrA variant lacking the C-terminal membrane anchor with His_6_-tagged Connectase from a Ni-NTA column (Fig. 1a). Surprisingly, this experiment also revealed the Thr-1 dependent formation of specific reaction products. Mass spectrometrical analysis identified these products as a C-terminal MtrA fragment (MtrA^C^; Fig. 1b) and a conjugate between the corresponding N-terminal MtrA fragment and Connectase (MtrA^N^-Connectase, Fig. 1c). This conjugate was 18 Da lighter than the combined mass of its components, indicating that it presents a covalent protein adduct formed by condensation.

**Fig. 1:**
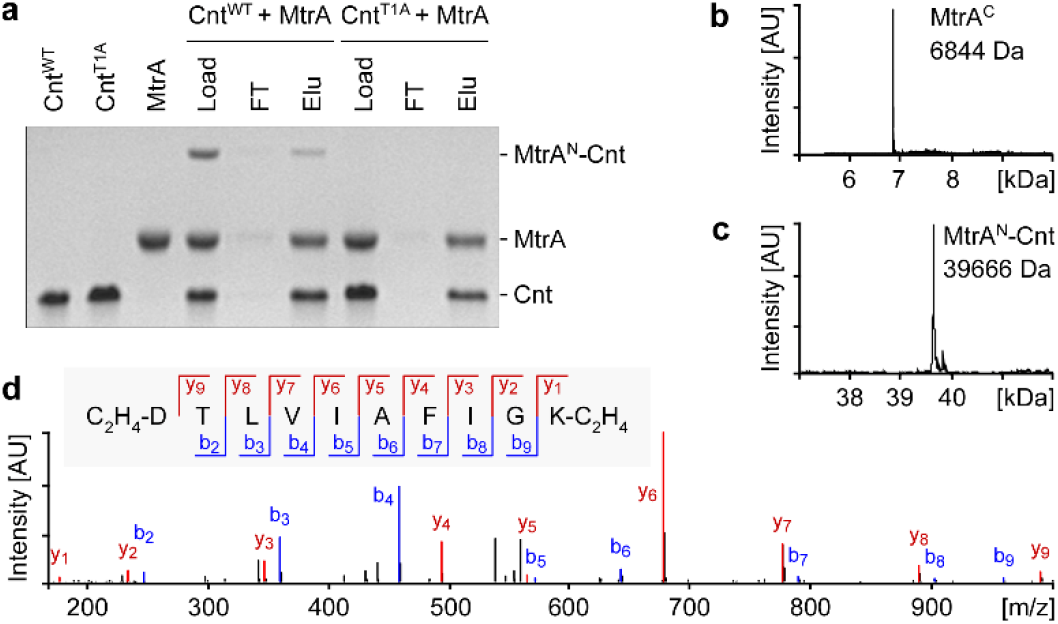
Connectase (Cnt) binds and modifies MtrA. (a) Polyacrylamide gel showing purified MtrA and His_6_-tagged Connectase (Cnt) proteins from *M. mazei*. When a mix of both (Load) is applied on a Ni-NTA column, MtrA is not found in the flow-through (FT) but co-elutes with Connectase (Elu). Furthermore, MtrA forms a specific reaction product (MtrA^N^-Connectase) with wild type Connectase (Cnt^WT^) but not with its active site mutant (Cnt^T1A^). (b, c) Newly generated masses in a *M. mazei* Connectase-MtrA reaction, as identified via liquid chromatography mass spectrometry (LCMS). These are interpreted as MtrA^C^ (residues 155-218; theoretical mass: 6588.4 Da / detected mass: 6844.7 Da) and as MtrA^N^-Connectase conjugate (39669.4 Da / 39665.8 Da). (d) MS/MS spectrum of the fusion peptide DTLVIAFIGK resulting from amide bond formation between the *M. mazei* Connectase N-terminus (TLVIAFIGK…) and the MtrA KDPGA-motif. The sample was digested with trypsin and free amino groups dimethylated (C_2_H_4_). The threonine amino group is not modified, suggesting its involvement in an amide bond.

The MtrA modification site contains a highly conserved KDPGA motif present in almost all Connectase encoding organisms (Extended Data, Fig. 4). Connectase processes MtrA between the aspartate and proline residues of this motif, resulting in MtrA^C^ (PGA…) and MtrA^N^-Connectase (…KD-Connectase). Accordingly, the tryptic fusion peptide (D-TLVIAFIGK) that results from a conjugation of MtrA with the Connectase N-terminus (TLVIAFIGK…) is abundant in each of the mass spectrometry datasets derived from pulldowns with whole cell extracts (see above). Moreover, when all free amino-groups of an MtrA-Connectase tryptic peptide mixture were methylated (dimethyl labeling,^13^), modifications in the above fragment were found only at the newly generated aspartate N-terminus and the lysine (Fig. 1d). By contrast, no methylation could be detected at the threonine, indicating that its amino group is engaged in a regular amide bond with the MtrA aspartate (Extended Data, Fig. 3b).

## Connectase recombines proteins via the MtrA (5)KDPGA(10) motif

To study the structural basis for this reaction, we conducted experiments with the Connectase^S1A^ active site mutant and MtrA from the thermophilic *M. jannaschii*, as *M. mazei* MtrA proved unsuitable for high-resolution analyses. Light scattering experiments show the monomeric nature of both proteins and their tight interaction within the heterodimer (Fig. 2a; Extended Data, Fig. 5a-b). Microscale thermophoresis (MST) experiments with synthetic peptides consisting of the KDPGA motif plus × N-terminal and y C-terminal residues from the MtrA sequence, denoted as (x)KDPGA(y) in the following, show that the (0)KDPGA(10) sequence is sufficient for this high-affinity interaction (Fig. 2b; Extended Data, Fig. 5e-l).

**Fig. 2:**
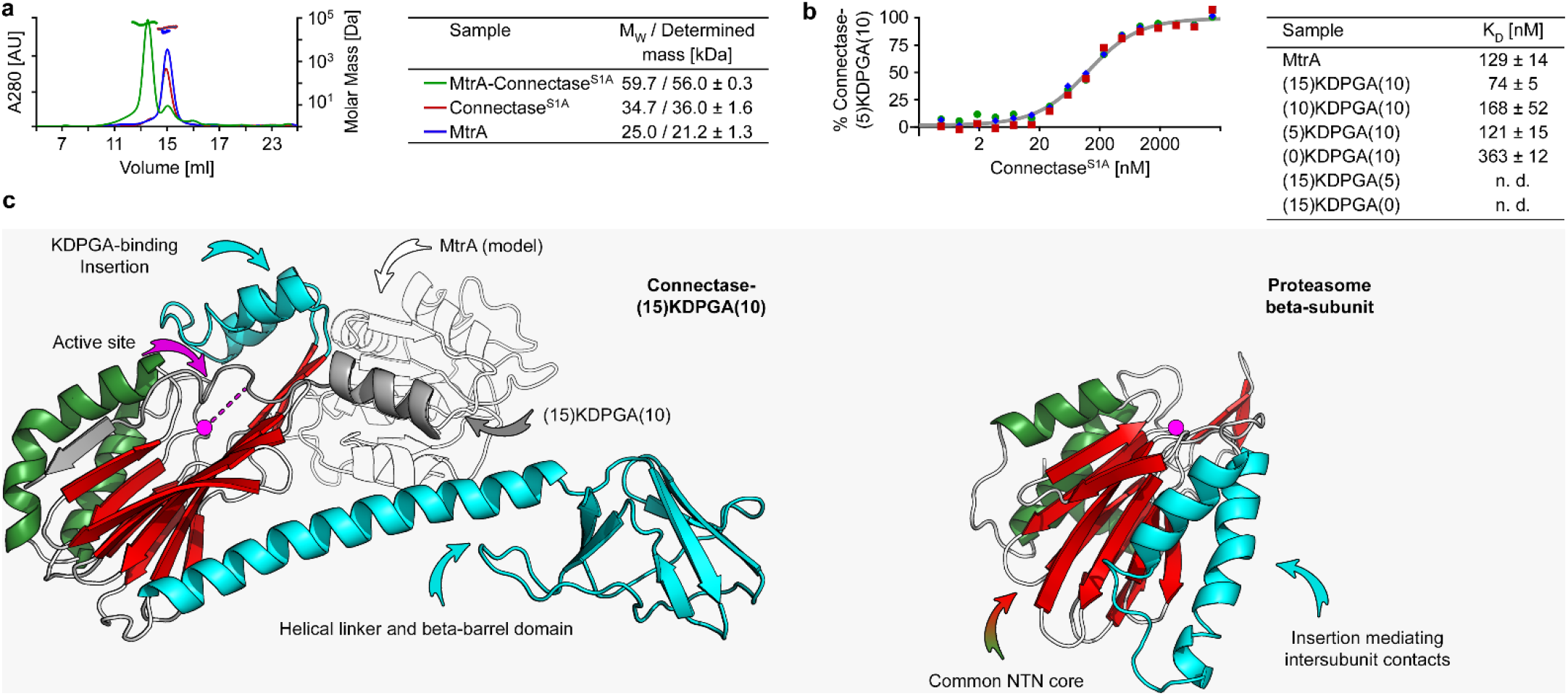
MtrA interacts with Connectase via a short amino acid motif, forming a heterodimer. (A) Gel filtration and light scattering analyses of *M. jannaschii* MtrA and Connectase^S1A^ proteins. While Connectase and MtrA alone show a comparable elution behavior, the mixture of both elutes at a lower volume, indicating complex formation (thin lines, plotted on the primary Y-axis). This interpretation is supported by light scattering measurements (thick lines, secondary Y-axis). The determined masses (table, right) closely resemble the theoretical monomeric masses for Connectase and MtrA alone and for the MtrA-Connectase heterodimer. (B) Binding curve visualizing the formation of a complex between the MtrA-derived peptide (5)KDPGA(10) and *M. jannaschii* Connectase^S1A^, as determined by microscale thermophoresis (MST). Analogous experiments with other substrates (table, right) show that the 15 amino acid peptide (0)KDPGA(10) is sufficient for this high affinity interaction. (C) Crystal structures of *M. jannaschii* Connectase^S1A^ in complex with (15)KDPGA(10) (left) and the proteasome beta subunit (right, PDB 3H4P^15^). Both proteins share a common NTN core domain (red and green) and an N-terminal active site residue (pink) but diverge in three protein-specific elements (cyan). A model based on an alignment with (15)KDPGA(10) shows, how MtrA (colorless, PDB 5L8X^12^) could potentially bind to Connectase.

We crystallized both Connectase alone and a (15)KDPGA(10)-Connectase^S1A^ complex and could solve their structures at 2.3 Å and 3.05 Å, respectively (Fig. 2c; Extended Data, Fig. 5m). Compared to proteasome subunits, Connectase lacks a two-helix element mediating inter-subunit contacts within the proteasome complex but features a two-helix insertion at a different position, which interacts with the KDPGA motif and might control substrate specificity. The C-terminal beta-barrel domain, which is found only in some Connectase variants, is connected via a long helical linker and may assist in binding of MtrA (Extended Data, Fig. 5c-d), but is generally not required for the Connectase reaction (see below). In the NTN domain, substrate binding is mediated via induced beta-sheet interactions with the KDPGA(10) sequence, locating the particularly fragile^14^ Asp-Pro bond to the active site cleft. In contrast, the helix preceding the KDPGA(10) motif is not bound specifically.

To verify these results, we incorporated the *M. mazei* MtrA-derived (5)KDPGA(10) motif in a different protein, *Caldiarchaeum subterraneum* ubiquitin^16^, and found that Connectase modifies it just like MtrA. Surprisingly though, when we followed the time course of the reaction, we found only a constant fraction of Ub-(5)KDPGA(10) modified at any time (Fig. 3b, lanes 1-4). This suggested a reversible reaction, resulting in an equilibrium between modified and unmodified Ub-(5)KDPGA(10) (Fig. 3a). In this scenario, the “forward” reaction yields Ub-(5)KD-Connectase and PGA(10), while the “reverse” reaction restores Ub-(5)KDPGA(10). To prove this assumption, we designed alternative substrates that could be used in the “reverse” reaction, PGA(10)-sdAb (single domain antibody), PGA(10)-CyP (Cyclophilin) and PGA(10)-GST (Glutathione-S-Transferase). These substrates could be conveniently produced via expression of MPGA(10)-sdAb/Cyc/GST in *E. coli*, following “automatic” methionine removal by the endogenous methionine aminopeptidase^17^. Indeed, a reaction with Ub-(5)KDPGA(10) and PGA(10)-sdAb/Cyc/GST resulted in equimolar ratios of educts and Ub-(5)KDPGA(10)-sdAb/Cyc/GST products (Fig. 3b, lanes 5-10; Extended Data, Fig. 6). Hence, Connectase is capable of ligating any two substrates featuring the (5)KDPGA(10) and PGA(10) motif, respectively.

**Fig. 3:**
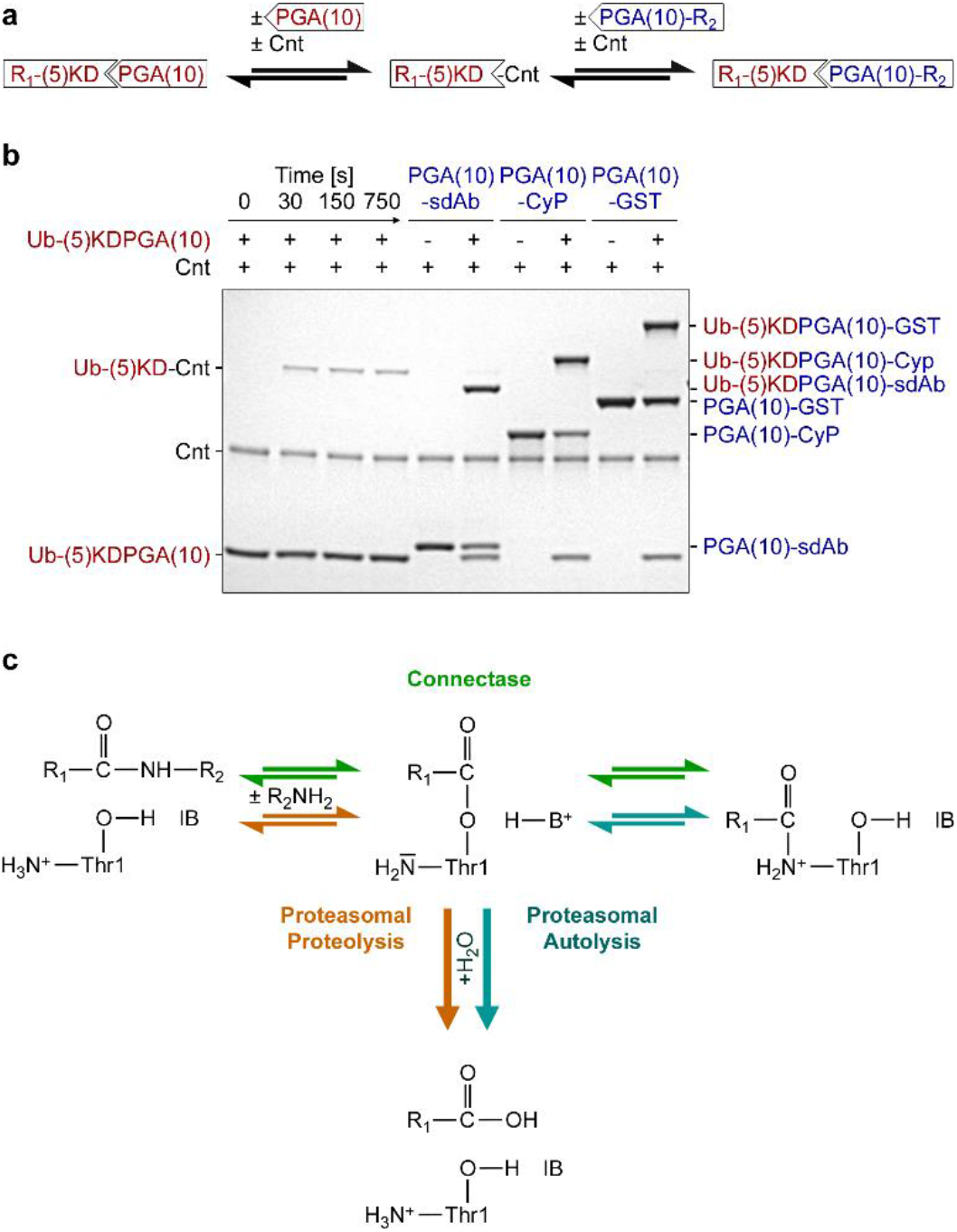
Connectase (Cnt) modifications are reversible and allow the recombination of substrates. (A) Schematic representation of an Connectase-mediated ligation of two protein substrates, R_1_-(5)KDPGA(10)-R_2_ and PGA(10)-R_3_. (B) Polyacrylamide gel showing the incubation of *M. mazei* Connectase with various proteins bearing the Connectase recognition motif. A constant fraction of Ub-(5)KDPGA(10) forms a conjugate with Connectase (Ub-(5)KD-Connectase), suggesting a reversible reaction between the two (lanes 1-4). In place of PGA(10), PGA(10)-sdAb/CyP/GST can be used as alternative substrates for the “reverse reaction”, resulting in an equilibrium of the respective educts and Ub-(5)KDPGA(10)-sdAb/CyP/GST products (lanes 5-10). (C) The proposed Connectase reaction mechanism (green) as a combination of the first steps of two known proteasomal reactions: proteolysis (orange) and autolysis (i.e. propeptide processing, cyan). A substrate, R_1_-R_2_ (left), is cleaved (middle) and the N-terminal fragment R_1_ transferred on the Connectase Thr-1/Ser-1 N-terminus (right; see Fig. 1D). The reaction is reversible, allowing substrate recombination with an alternative R_2_ substrate (see Fig. 3A). It differs from proteolysis/autolysis by avoiding the irreversible hydrolysis step (bottom).

Based on the reversibility of the reaction, the identification of an amide-bonded intermediate (Fig. 1d) and the requirement of an active-site hydroxyl group as the catalytic nucleophile (Fig. 1a), we propose that the Connectase catalytic mechanism is a variation of steps involved in two known proteasomal reactions, hydrolysis and autolysis^18^, but differs from them by avoiding the final irreversible hydrolysis step (Fig. 3c). The underlying reaction sequence is known as ordered ping-pong mechanism, meaning that the reaction can only start with the primary R_1_-(5)KDPGA(10) substrate and only proceed with the secondary PGA(10)-R_2_ substrate (R_1/2_ = any molecule). As both substrates utilize the same binding site, reaction rate and product yield are influenced by primary/secondary substrate ratios. In this model, the highest molar product/educt1+educt2 yield can be achieved with equimolar substrate ratios, making this the set-up of choice for most protein-protein ligations. Nevertheless, the use of different substrate ratios can be useful, where complete modification of one reaction partner is desired, e.g. when labeling a protein with a fluorophore.

## Connectase catalyzes specific and efficient ligations without side reactions

A more detailed characterization of the reaction shows that Connectase-catalyzed protein-protein ligations at a molar enzyme:substrate ratio of 1:400 usually result in an equilibrium of equimolar substrates and products within minutes (Fig. 4a). A systematic analysis of sequence determinants N- and C-terminally of the MtrA-derived (x)KDGPA(y) / PGA(z) motif suggests that 5 residues N-terminally of KDPGA and 10 residues C-terminally of KDPGA / PGA allow efficient ligations in most cases, but that sterically demanding protein-protein ligations are much faster with 15 residues present C-terminally of PGA (Fig. 4b, Extended Data, Fig. 7). The maximum rate of such an exemplary *M. mazei* Connectase catalyzed protein-protein ligation using Ub-(5)KDPGA(15) and PGA(15)-Ub was determined with k_cat_ = 0.92 ligations per enzyme and second. The half-maximum reaction speed was observed at a substrate concentration of ~2.2 μM each (Fig. 4c; Extended Data, Fig. 8). Similar ligation rates were found in reactions with full-length *M. jannaschii* Connectase, while a deletion of the beta-barrel domain led to a slight decrease in reactivity (Extended Data, Fig. 9).

**Fig. 4:**
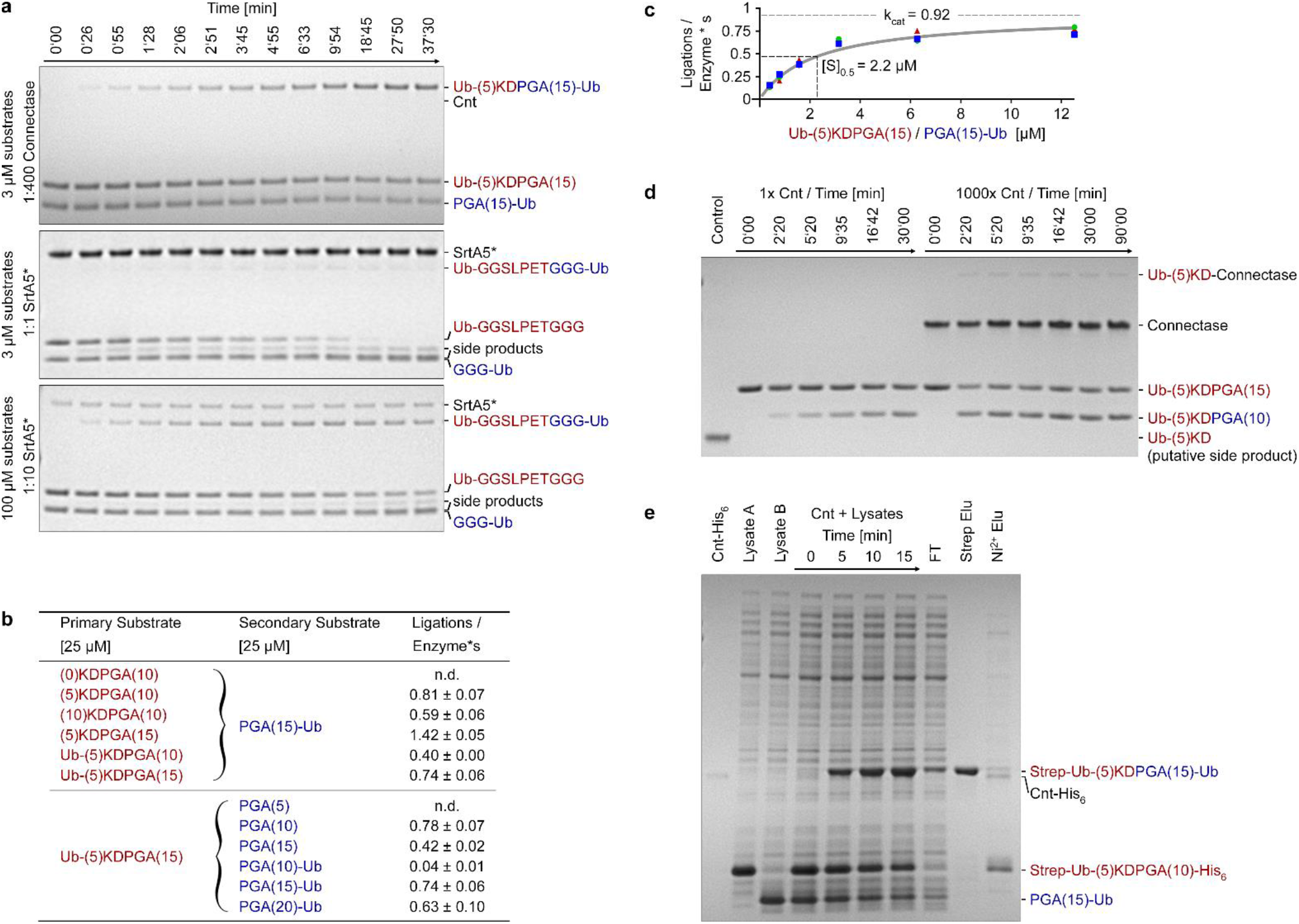
Connectase (Cnt) catalyzes specific and efficient ligations without side reactions. (A) The time course of *M. mazei* Connectase and Sortase A pentamutant (SrtA5*)-mediated ligations of two ubiquitin molecules at various molar enzyme:substrate (1:1, 1:10, 1:400) concentrations. Connectase is used at very low concentrations and therefore not visible on the gel. (B) Ligation rates for Connectase reactions with various peptide and protein substrate pairs at 25 μM. (C) Michaelis-Menten plot of a *M. mazei* Connectase ligation using Ub-(5)KDPGA(15) and PGA(15)-Ub substrates. Although the Connectase reaction can be expected to follow more complex mathematics, the experimental data is well described by the Michaelis-Menten model (grey line). (D) Polyacrylamide gel showing a test for *M. mazei* Connectase hydrolase activity with Ub-(5)KDPGA(15) and PGA(10) substrates. At an enzyme:substrate concentration of 1:1000 (1x), Connectase catalyzes ~50% product formation (i.e. Ub-(5)KDPGA(10)) in 30 minutes without forming the putative hydolysis product Ub-(5)KD. Even at 1000x increased enzyme concentrations and prolonged incubation times, no hydrolysis product can be detected. (E) Polyacrylamide gel showing the ligation of two recombinantly expressed substrates (Strep-Ub-(5)KDPGA(10)-His_6_ and PGA(15)-Ub) in cell lysates and the single step purification of the respective ligation product (Strep-Ub-(5)KDPGA(15)-Ub) using a Ni-NTA column in series with a Streptavidin column.

We then sought to compare these rates with the most widely used enzyme ligase, Sortase A (SrtA), which has proven a powerful and reliable tool in numerous remarkable applications^19–22^. SrtA has been extensively optimized^23,24^ and state of the art in many labs is a SrtA pentamutant (SrtA5*) with significantly increased activity^25^. Analogous to Connectase, SrtA ligates two sequences bearing the LPET-G motif and an N-terminal glycine, respectively, though additional linker sequences are usually introduced to avoid steric hindrances and to increase reactivity^7,26,27^ In addition, SrtA also catalyzes the irreversible hydrolysis of substrates and products featuring this motif at a lower rate (k_cat Ligation_/k_cat Hydrolysis_ ≈ 3.3^28^) as well as side reactions, in which lysine side-chains substitute the N-terminal glycine substrate. The maximum amount of product can therefore only be obtained by monitoring the ligation/hydrolysis ratio and by stopping the reaction at just the right time. Following established protocols^26,29^, we designed Ub-GGSLPETGGG and GGG-Ub substrates and recorded the time course of the reaction. In our assays, SrtA5* displayed ~100x higher ligase activity compared to SrtA, but also catalyzed more side reactions, possibly due to its low affinity for the secondary substrate (K_M LPETG_ = 170 μM; K_M GGG_ = 4700 μM^25^; Extended Data, Fig. 10). Unlike Connectase, both Sortase variants showed only spurious ligase activity at low substrate concentrations (3 μM each), but displayed far more favorable characteristics at high substrate concentrations (100 μM each; Fig. 4a and Extended Data, Fig. 10). Yet, even under those conditions, Connectase shows >4000x higher ligase activity than non-optimized SrtA and >40x increased ligase activity compared to optimized SrtA5*. Furthermore, we observed increased ligation yields in Connectase reactions (~50% compared to ~30%), which we attribute to the apparent absence hydrolysis side reactions. These results are comparable with Sortase ligations of other protein substrates^7,29,30^ and demonstrate that Sortase is a good choice where short ligation motifs are preferred and high substrate concentrations available^31^. Where these requirements are not met, Connectase is advantageous, as it combines substantially higher substrate affinities, reaction rates and ligation yields without catalyzing detectable side reactions.

To study whether Connectase catalyzes such side reactions under more extreme conditions, we extended incubation times to several hours and increased Connectase concentrations by 1000x (Fig. 4d). Over the entire period, the amount of ligation product remained constant at ~50%, while no extra products could be detected. This finding may surprise, considering that all currently known ligase enzymes also function as proteases by forming an ester-bond with the substrate that is then either transferred to water (hydrolysis/proteolysis) or to the amino group of a ligation partner^6^. Connectase, however, differs from these enzymes in using the NTN active site architecture that allows a third alternative: the transfer to the N-terminal amino-group and the associated formation of a hydrolysis resistant amide-bonded intermediate (Fig. 3c).

To test whether the above characteristics hold true in more complex solutions, we ligated two independently expressed protein substrates, Strep-Ub-(5)KDPGA(10)-His_6_ and PGA(15)-Ub, within their respective cell lysate and subsequently purified the ligation products in a single step, using a Ni-NTA column in series with a streptavidin column (Fig. 4e). Only one protein species, Strep-Ub-(5)KDPGA(15), could be eluted with desthiobiotin, indicating that no other proteins in the lysate reacted with the Strep-Ub-(5)KD-Connectase intermediate. In accordance with the nanomolar affinity interaction between Connectase and its conserved recognition motif (Fig. 2b), this observation suggests that Connectase ligations are highly specific and applicable even in complex solutions. Moreover, this experiment highlights the feasibility of Connectase-mediated ligations for the large-scale generation and single-step purification of a given ligation product in short time and with minimal amounts of enzyme.

## Conclusion

Connectase is a favorable new enzyme for protein conjugation. As such, it complements existing methods, such as native chemical ligation^1^, expressed protein ligation^2^, enzyme-mediated ligations^5^, protein trans-splicing^3,32^ or split domain systems^4^. All of these methods are widely used, each with its functional niche due to its respective advantages and limitations^6,33–36^. While Connectase ligations require a longer linker between the ligated substrates than some of these methods, it also offers a unique combination of simplicity and efficiency: robust, fast and highly specific ligations without side reactions that require no lengthy optimization procedures, complicated set-up, special equipment or defined buffer systems. Substrates can be produced recombinantly, but are also amenable to chemical synthesis. With these characteristics, Connectase is highly attractive for a variety of applications, ranging from protein immobilization and fusion to chemical compounds (e.g. fluorophores) to protein-protein ligations.

## Acknowledgments

We thank Anja Rau, Lorena Vöhringer and Eva Hertle (MPI for Developmental Biology) for technical assistance, Ruth Schmitz-Streit and Katrin Weidenbach (University of Kiel) for providing *Methanosarcina mazei* cell extract, and Victoria Sanchez (Core facility, MPI for Biochemistry, Martinsried) as well as Mirita Franz-Wachtel (Proteome Center, University of Tübingen) for mass spectrometric analyses. We thank Kerstin Bär and Reinhard Albrecht for assistance with crystallographic sample preparation, Reinhard Albrecht and Christopher Heim for crystallographic data collection, and the staff of beamline X10SA/Swiss Light Source for excellent technical support. This work was supported by institutional funds from the Max Planck Society.

## Data availability

Protein structure coordinates and structure factors have been deposited in the Protein Data Bank under accession codes 6ZVZ (wild-type Connectase) and 6ZW0 (complex of the Connectase S1A mutant with peptide). Nucleotide and amino acid sequences are provided in the Supplementary Information. All relevant data are available from the corresponding author upon request.

## Materials & Correspondence

All correspondence and material requests should be addressed to Andrei N. Lupas.

## Author contributions

MA discovered the homology of DUF2121 to proteasome subunits and initiated the project. ACDF discovered the ligase activity of DUF2121 and performed most experiments reported here. MDH solved the crystal structures. MDH, JM and ANL supervised the project and discussed all results. ACDF wrote the paper with input from all authors.

