## Extended Data for "The proteasome-related Connectase is a fast and specific protein ligase"

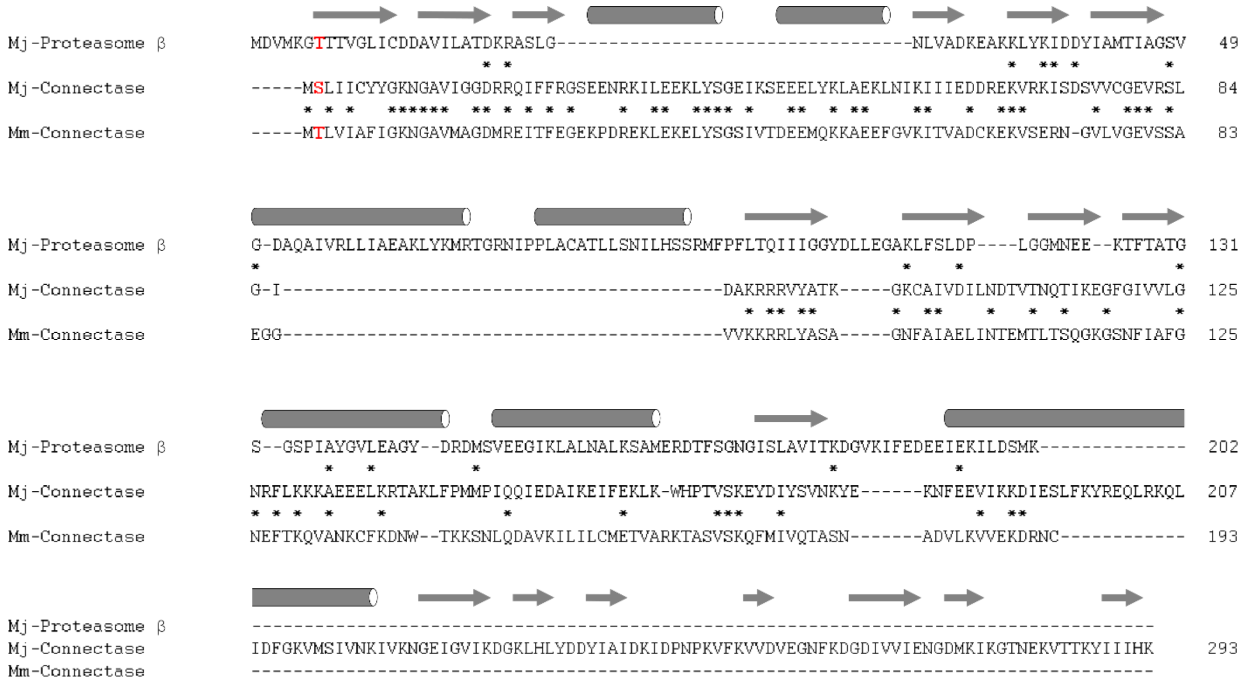


**Extended Data, Figure 1: Connectase is a distant proteasome homolog with distinct structural features**

Structure-based sequence alignment of the *Methanocaldococcus jannaschii* proteasome β subunit, *M. jannaschii* Connectase and *Methanosarcina mazei* Connectase. Secondary structure elements are indicated by arrows (beta-sheets) and tubes (helices). Identical residues between two sequences are marked by asterisks and catalytic Thr / Ser residues are shown in red. The alignment visualizes the distant relationship between proteasomal and Connectase NTN (N-terminal nucleophile) domains. Compared to proteasome subunits, Connectase lacks a two-helix element mediating inter-subunit contacts within the proteasome complex but features a two-helix insertion at a different position. Some Connectase variants, such as the one from *M. jannaschii*, contain an additional C-terminal six-stranded beta-barrel domain that is connected to the NTN domain through a long helix. The propeptide present in most proteasome beta subunits is absent from Connectase, whose start-methionine is most likely removed by endogenous methionine aminopeptidases. Both proteins share conserved active site residues.

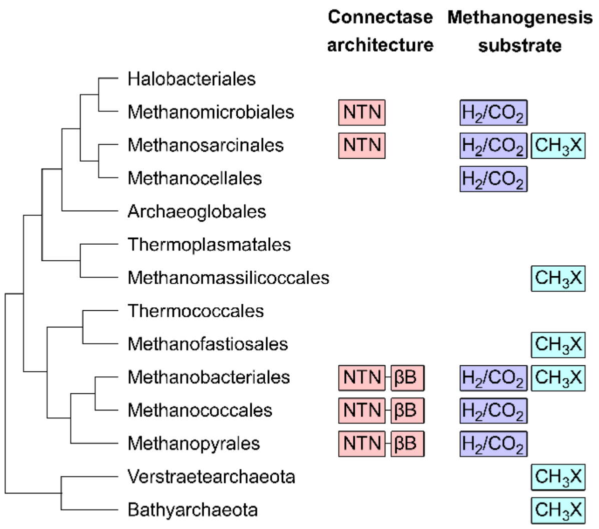


**Extended Data, Figure 2: The distribution of Connectase proteins is linked to hydrogenotrophic methanogenesis.**

Shown is the phylogenetic distribution of Connectase proteins in methanogenic archaea and related orders. While being composed of just the regular proteasome-type NTN-domain (N-terminal nucleophile) in class II methanogens [1], Connectase from class I methanogens features an extra C-terminal beta-barrel domain (βB). Both variants almost always co-occur with genes involved in methanogenesis from carbon dioxide and molecular hydrogen (hydrogenotrophic methanogenesis [2]). By contrast, their distribution is not linked to methanogenesis from other substrates, such as acetate, methyl amine or methanol ($CH_3X$).

**a**

| Rank by Intensity | Detected Protein | Intensity (Experiment 1) | Intensity (Experiment 2) | Intensity (Experiment 3) |
|---|---|---|---|---|
| 1 | Connectase | 10000 | 10000 | 10000 |
| 2 | MtrA | 50 | 721 | 1952 |
| 26 | MtrH | 94 | 96 | 95 |
| 263 | MtrG | 1 | 2 | 7 |
| 325 | MtrB | 2 | 5 | 3 |
| 373 | MtrE | 3 | 3 | 2 |
| 661 | MtrF | 0 | 0 | 0 |

**b**

| Fragment | Sequence [* = $C_2H_6$ modification] | Intensity L ($MtrA^N$-Connectase) | Intensity M (MtrA) | Intensity H (Connectase) |
|---|---|---|---|---|
| MtrA 125-146 | *FQEQVQVVNLLDTEDMGAITSK* | 496 | 1137 | 0 |
| MtrA 154-191 | *DPGAFDADPLVVEISEEGEEEEEGGVVRPVSGEIAVLR | 0 | 100 | 0 |
| MtrA 201-209 | *MMDIGNLNK* | 53 | 10000 | 1 |
| MtrA 149-154 + Connectase 1-9 (amide) | *ELASKDTLVIAFIGK* | 48 | 0 | 0 |
| MtrA 149-154 + Connectase 1-9 (ester) | *ELASKD*TLVIAFIGK* | 0 | 0 | 0 |
| MtrA D154 + Connectase 1-9 (amide) | *DTLVIAFIGK* | 1975 | 0 | 0 |
| MtrA D154 + Connectase 1-9 (ester) | *D*TLVIAFIGK* | 0 | 0 | 0 |
| Connectase 10-19 | *NGAVMAGDMR | 407 | 0 | 1723 |
| Connectase 1-9 | *TLVIAFIGK* | 85 | 1 | 1639 |

**Extended Data, Figure 3 (related to Fig. 1): The Connectase N-terminus forms an amide bond with MtrA D154.**

(a) Mass spectrometrically detected proteins in pulldowns with *M. mazei* Connectase coupled to Strep- (Experiment 1), Myc- (Experiment 2) or HA-tags (Experiment 3) as bait, using *M. mazei* whole cell extract. MtrA is detected at high intensities (normalized and non-quantitative), while a weaker signal is determined for other subunits of the MtrA-MtrH complex.

(b) Detected protein fragments in excised polyacrylamide gel bands corresponding to Connectase (H), MtrA (M) or an $MtrA^N$-Connectase conjugate (L; see Fig. 1a). The samples were digested with trypsin and dimethylated at primary amine groups (indicated by asterisks), using different isotopes (H = Heavy; M = Medium; L = Light). Note, that the relative intensities (normalized to 10000) for a given peptide reflect quantitative differences between the samples. The band corresponding to $MtrA^N$-Connectase also contains small amounts of unconjugated MtrA and Connectase proteins, possibly due to the reversibility of the reaction.

| | |
|---|---|
| Sequence Logo | 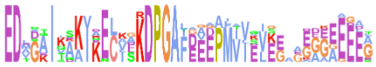 |
| *Methanofollis ethanolicus* | (138)EDLGAIKAKIAELKGKDPGAFGEAPIVVEVKEEGG(84) |
| *Methanoculleus marisnigri* | (138)EDMGAIKAKIDELKARDPGAFGAEPMIVEVKEAGG(82) |
| *Methanolinea tarda* | (138)EDLGAIKAKIAELKGRDPGAFAADPMVVEVKEAAG(83) |
| *Methanopyrus kandleri* | (147)EDVDEIVKAIEECVEKDPGAYEEGPMTISLEEEEE(85) |
| *Methanobrevibacter smithii* | (137)EDVGAIQAKINECVEKDPGAFEEEAMVISVEGDDG(81) |
| *Methanobacterium formicicum* | (139)EDAATIQSKVKECIDKDPGAFEEEAMVIVVEEDGE(80) |
| *Methanothermobacter thermautotrophicus* | (139)EDADAIKAKVKECIEKDPGAFEEEAMVIKVEEGGE(79) |
| *Methanothermus fervidus* | (138)EDADQIKEKVKECIEKDPGAFEEEAMVVKVEEEEE(79) |
| *Methanothrix thermacetophila* | (139)EDMGAITAKIKELVAKDPGALDVEPMIVEIKEGAG(84) |
| *Candidatus Methanoperedens BLZ2* | (138)EDENAIIAKIKELAAKDPGAFDGEPMIIQVGEAKE(83) |
| *Methanosarcina mazei* | (138)EDMGAITSKVRELASKDPGAFDAEPLVVEISEEGE(82) |
| *Methanosaeta harundinacea* | (139)EDTGTIVSKIKECVSKDPGALDVEPMAVEISEGEA(83) |
| *Methanococcus maripaludis* | (140)EDTGKISDAIKNCISKDPGAFEEEPMVIELEGGAA(79) |
| *Methanocaldococcus jannaschii* | (144)EDIGKITQAIKECLSKDPGAIDEDPFIIELEGGKG(81) |
| *Methanotorris igneus* | (140)EDVSKITAAIKECISKDPGAIDEEPMVLDLEGGGA(83) |

**Extended Data, Figure 4: The MtrA KDPGA motif is highly conserved.**

An alignment of the KDPGA motif with the 15 preceding and following amino acids ((15)KDPGA(15)) in a phylogenetically diverse set of MtrA proteins. The sequence conservation is visualized above the alignment, with larger letters indicating higher conservation. All shown MtrA proteins co-occur with Connectase.

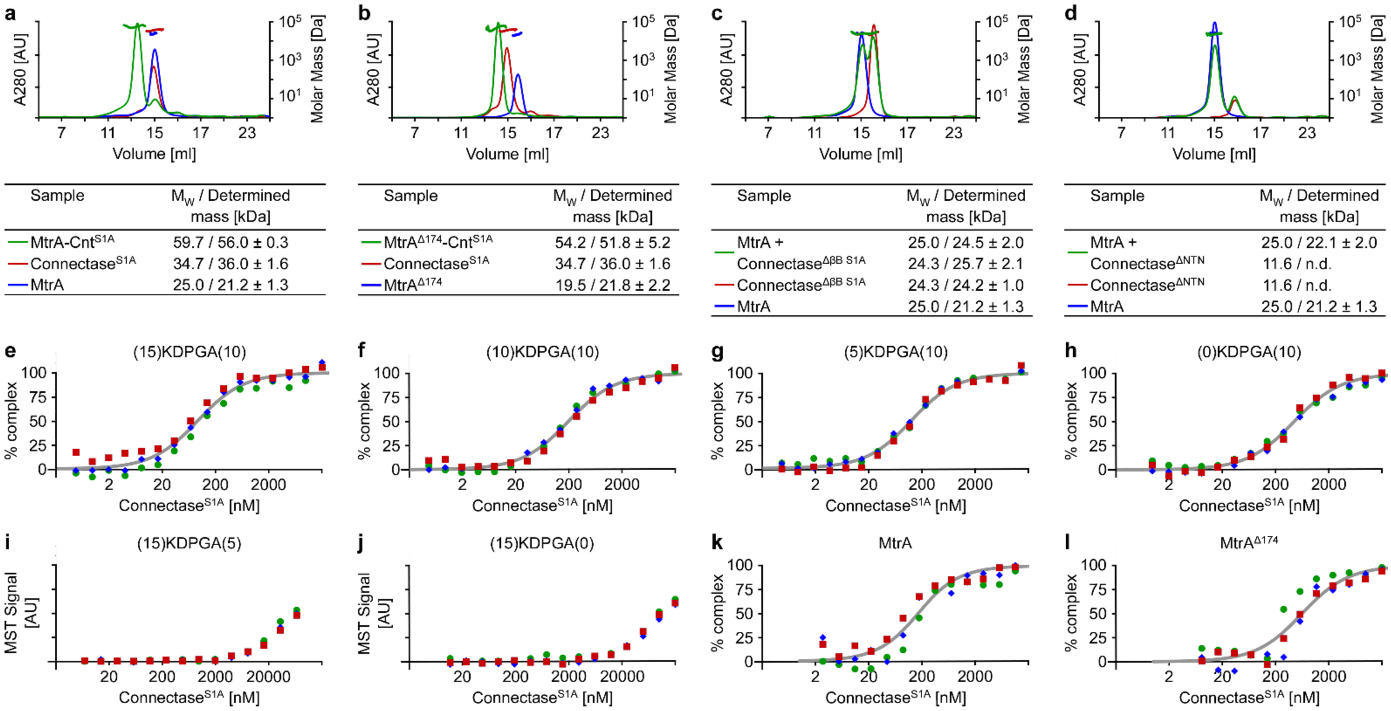


m

| | Connectase | | Connectase [Se-Met] | | | $Connectase^{S1A}$-(15)KDPGA(10) |
|---|---|---|---|---|---|---|
| **Data collection** | | | | | | |
| Space group | C2 | | C2 | | | $P2_12_12_1$ |
| Cell dimensions | | | | | | |
| a, b, c (Å) | 76.2, 190.2, 111.9 | | 76.2, 192.1, 113.0 | | | 72.2, 98.5, 109.5 |
| α, β, γ (°) | 90.0, 107.4, 90.0 | | 90.0, 107.6, 90.0 | | | 90.0, 90.0, 90.0 |
| | | *Low Remote* | *Inflection* | *Peak* | *High Remote* | |
| Wavelength (Å) | 1.000 | 0.9863 | 0.9790 | 0.9785 | 0.9709 | 1.000 |
| Resolution (Å) | 37.2 – 2.30 | 38.1 – 3.00 | 38.1 – 3.00 | 40.0 – 2.80 | 38.1 – 3.00 | 49.3 – 3.05 |
| | (2.44 – 2.30) | (3.18 – 3.00) | (3.18 – 3.00) | (2.97 – 2.80) | (3.18 – 3.00) | (3.24 – 3.05) |
| Rsym or Rmerge | 7.7 (74.7) | 8.3 (53.6) | 10.3 (64.9) | 7.3 (64.7) | 9.4 (60.7) | 14.4 (99.3) |
| *I* / σ*I* | 11.6 (1.69) | 11.5 (2.01) | 9.32 (1.63) | 12.3 (1.68) | 10.2 (1.77) | 10.5 (1.81) |
| Completeness (%) | 97.7 (95.8) | 99.4 (98.1) | 99.4 (98.1) | 99.5 (98.8) | 99.5 (98.7) | 99.6 (99.4) |
| Redundancy | 4.31 (4.30) | 3.52 (3.56) | 35.2 (35.6) | 3.51 (34.8) | 3.51 (3.54) | 6.16 (5.89) |
| **Refinement** | | | | | | |
| Resolution (Å) | 37.2 – 2.30 | | | | | 49.3 – 3.05 |
| No. reflections | 65793 | | | | | 14598 |
| $R_{work}$ / $R_{free}$ | 24.2 / 27.8 | | | | | 25.4 / 29.4 |
| No. atoms | 9599 | | | | | 5322 |
| Protein | 9554 | | | | | 5264 |
| Ligand/ion | - | | | | | 58 |
| Water | 45 | | | | | - |
| B-factors | 64.8 | | | | | 93.5 |
| Protein | 65.0 | | | | | 93.4 |
| Ligand/ion | - | | | | | 98.0 |
| Water | 42.0 | | | | | - |
| R.m.s deviations | | | | | | |
| Bond lengths (Å) | 0.014 | | | | | 0.004 |
| Bond angles (°) | 1.55 | | | | | 0.87 |

**Extended Data, Figure 5 (related to Fig. 2): MtrA interacts with Connectase (Cnt) via a short amino acid motif, forming a heterodimer**

(a) Gel filtration and light scattering analyses of *M. jannaschii* MtrA and $Connectase^{S1A}$ proteins, as shown in Fig. 1A. While Connectase and MtrA alone show a comparable elution behavior, the mixture of both elutes at a lower volume, indicating complex formation (elution profiles plotted on the primary Y-axis). This interpretation is supported by light scattering measurements analysing the molecular mass of proteins in the respective peaks (secondary Y-axis and table).

(b-d) Analogous experiments using alternative MtrA or Connectase substrates. MtrA-Connectase co-elution is still observed when residues C-terminal of the MtrA KDPGA(10) motif are deleted (b, $MtrA^{\Delta 174}$), but not with Connectase variants lacking either the C-terminal beta-barrel domain (c, $Connectase^{\Delta \beta B\ S1A}$) or the NTN-domain (d, $Connectase^{\Delta NTN}$). While these results suggest that the C-terminal beta-barrel domain assists in binding of MtrA, its presence is not required for Connectase-mediated ligations (Extended Data, Fig. 9).

(e-l) MST measurements for determination of the $K_D$-values in Fig. 1b.

(m) Data collection, phasing and refinement statistics for crystal structures. Values in parentheses are for the highest-resolution shell.

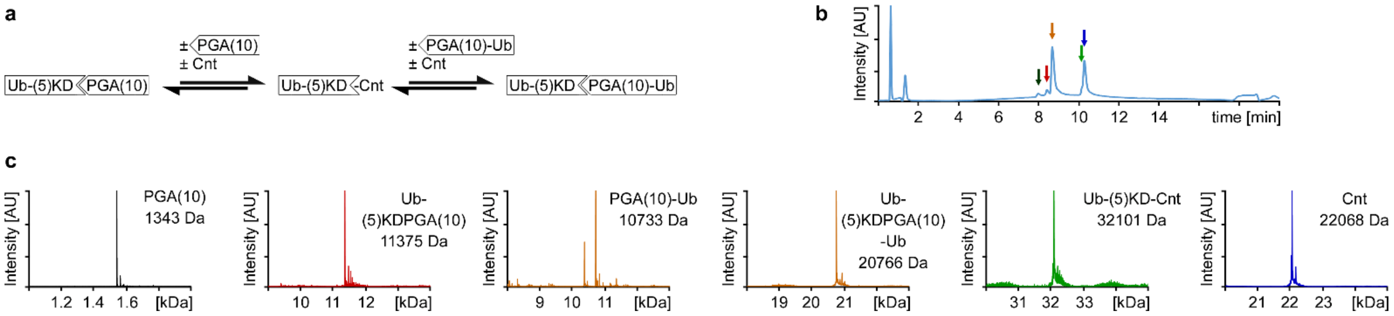


**Extended Data, Figure 6 (related to Fig. 3): Mass spectrometrical analysis of a Connectase-catalyzed protein-protein ligation.**

(A) Schematic representation of the Connectase reaction with Ub-(5)KDPGA(10) and PGA(10)-Ub substrates.

(B) Chromatographic elution profile of the same reaction.

(C) Mass spectrometrical analysis of the peaks determined in (B), i.e. 7.9 - 8.0 min (black graph), 8.3 - 8.5 min (red), 8.6 - 8.8 min (orange), 10.11 - 10.17 min (green) and 10.24 - 10.31 (blue). The masses of all products described in (A) are detected, while the mass of a potential hydrolysis side-product (i.e. Ub-(5)KD, 10.05 kDa) is not detected.

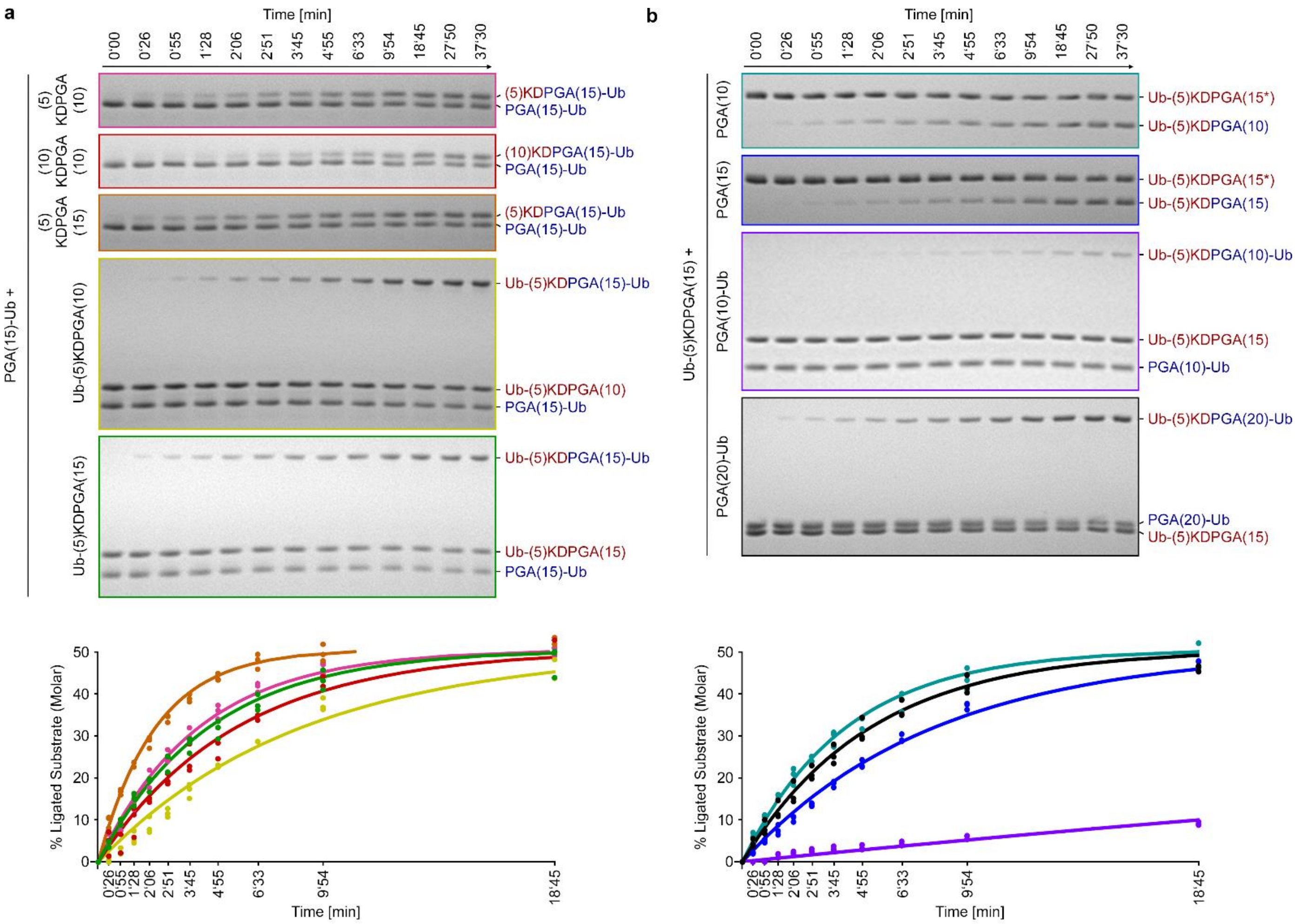


| Primary Substrate [25 µM] | Secondary Substrate [25 µM] | Ligations / Enzyme*s |
|---|---|---|
| (0)KDPGA(10) | PGA(15)-Ub | n.d. |
| (5)KDPGA(10) | PGA(15)-Ub | 0.81 ± 0.07 |
| (10)KDPGA(10) | PGA(15)-Ub | 0.59 ± 0.06 |
| (5)KDPGA(15) | PGA(15)-Ub | 1.42 ± 0.05 |
| Ub-(5)KDPGA(10) | PGA(15)-Ub | 0.40 ± 0.00 |
| Ub-(5)KDPGA(15) | PGA(15)-Ub | 0.74 ± 0.06 |

| Primary Substrate [25 µM] | Secondary Substrate [25 µM] | Ligations / Enzyme*s |
|---|---|---|
| Ub-(5)KDPGA(15) | PGA(5) | n.d. |
| Ub-(5)KDPGA(15) | PGA(10) | 0.78 ± 0.07 |
| Ub-(5)KDPGA(15) | PGA(15) | 0.42 ± 0.02 |
| Ub-(5)KDPGA(15) | PGA(10)-Ub | 0.04 ± 0.01 |
| Ub-(5)KDPGA(15) | PGA(15)-Ub | 0.74 ± 0.06 |
| Ub-(5)KDPGA(15) | PGA(20)-Ub | 0.63 ± 0.10 |

**Extended Data, Figure 7 (related to Fig. 4b): (5)KDPGA(10) is a good ligation motif for sterically accessible substrates, whereas (5)KDPGA(15) is efficient for sterically more demanding substrates.**

(a, b) Time courses of M. *mazei* Connectase-mediated ligations of various peptide or protein substrate pairs, visualized via SDS-PAGE and analyzed by band quantification (plots). Shown are sample gels representing three independent experiments. Substrates were used at 25 µM, in 400x molar excess over Connectase. For the ligation with Ub-(5)KDPGA(15) and PGA(10) / PGA(15) peptides, an additional Strep-tag sequence (indicated by an asterisk) was added C-terminal of the Ub-(5)KDPGA(15) substrate, so that the ligation results in a size shift. The data fits allow the determination of the rate coefficients depicted in tables. No Connectase activity was observed with (0)KDPGA(10) or PGA(5) substrates (n.d.).

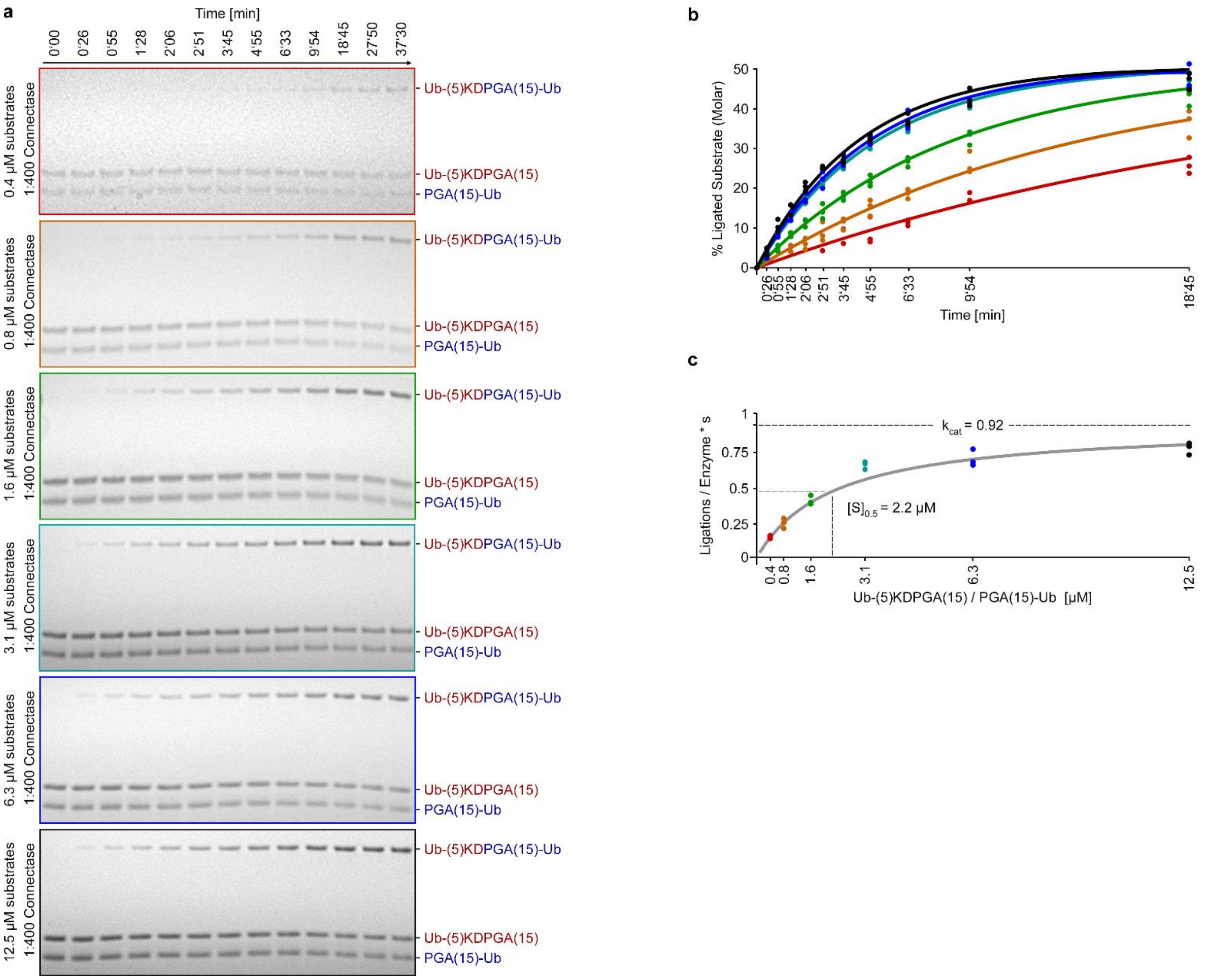


**Extended Data, Figure 8 (related to Fig. 4c): Kinetic analysis of a Connectase-catalyzed protein-protein ligation.**

(a) SDS-polyacrylamide sample gels depicting the time course of the same ligation reaction at various substrate concentrations (color coded).

(b) Quantification of ligation reactions, as shown in (a).

(c) Michaelis-Menten plot with ligation rate coefficients based on the data fits in (b). The kinetic parameters ($k_{cat}$ and $[S]_{0.5}$) are approximated and based on additional data (not shown) for higher substrate concentrations: 0.74 ± 0.05 $sec^{-1}$ at 25 µM, 0.88 ± 0.07 $sec^{-1}$ at 50 µM, and 0.90 ± 0.14 $sec^{-1}$ at 100 µM substrate concentration.

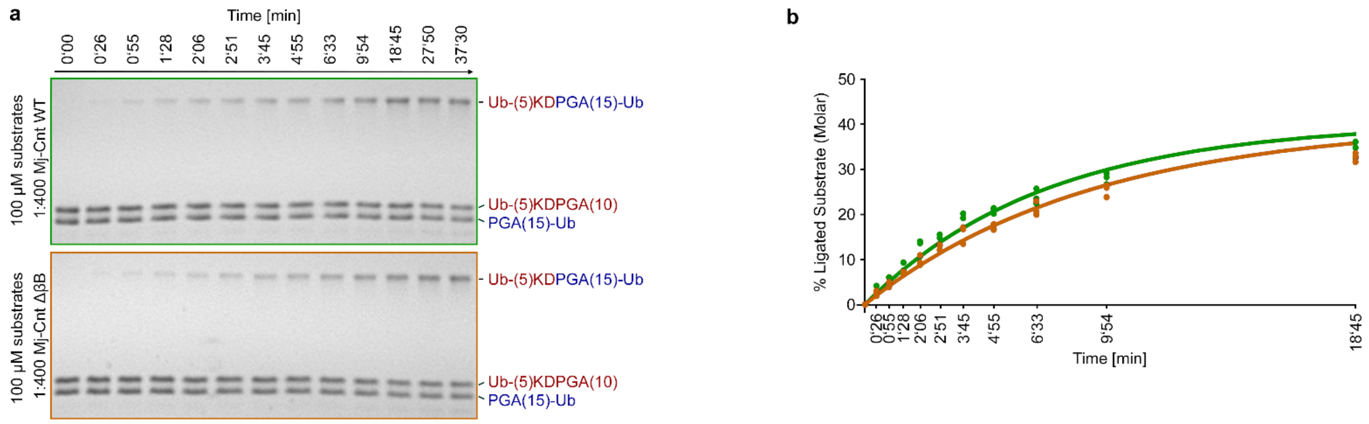


**Extended Data, Figure 9: The Connectase ligase reactivity is conserved in phylogenetically distant organisms and not dependent on the presence of the C-terminal beta-barrel domain.**

(a, b) SDS-polyacrylamide sample gels and derived plot showing the time course of *M. jannaschii* Connectase-catalyzed ligations, analogous to *M. mazei* Connectase reactions depicted in Extended Data, Fig. 7 and 8. Although both proteins share only 36% sequence identity, they catalyze protein-protein ligations at comparable rates, irrespective of the presence (green) or absence (orange) of the C-terminal beta-barrel domain.

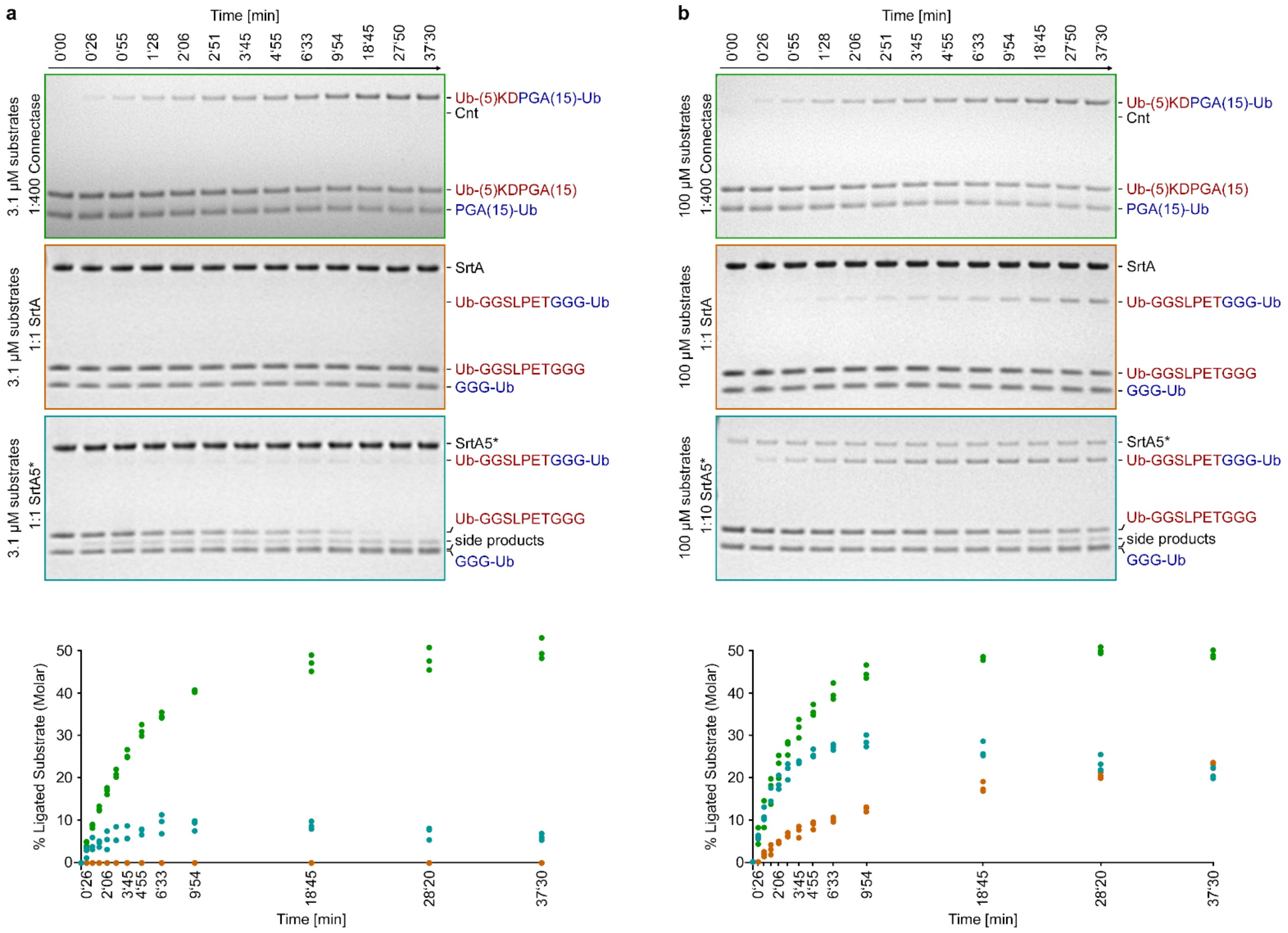


**Extended Data, Figure 10 (related to Fig. 4a): Comparison of Connectase- and Sortase-mediated protein-protein ligations.**

(a) SDS-polyacrylamide sample gels and derived plot showing the time course (triplicates) of comparable protein-protein ligations using *M. mazei* Connectase (green), *S. aureus* Sortase A (SrtA, orange) or evolved SrtA pentamutant (SrtA5*, cyan). Although Connectase is used at a 400x lower concentration, it catalyzes the ligation at much higher rates and without side-products.

(b) The same experiments at 32x higher substrate concentrations (i.e. 100 µM). Although these conditions are much more suitable for the Sortase reaction, Connectase shows ~4000x higher ligase activity compared to non-optimized (SrtA) and ~40x higher ligase activity compared to optimized (SrtA5*) Sortase enzymes.
