## Supplementary material for "The proteasome-related Connectase is a fast and specific protein ligase": Methods

***Bioinformatics***

Connectase as was identified through HHpred [1,2] searches with archaeal proteasome beta subunits against the Pfam database. The sequence alignment (Extended Data, Fig. 1) was generated with PromalS3D [3], based on the crystallographic structures of the *M. jannaschii* proteasome beta subunit (PDB 3H4P, [4]) and *M. jannaschii* Connectase (see crystallography section). The mapping of Connectase proteins on a phylogenetic tree (Extended Data, Fig. 2) was done based on previously described phylogenetic realationships and metabolic analyses [5,6].

***Cloning, Expression and Purification***

Connectase genes MM_2909 and MJ_0548 as well MtrA genes MM_1543 and MJ_0851 were amplified via PCR from genomic DNA (DSM3647 and DSM2661 (DSMZ)), cloned into pET30 vectors and expressed recombinantly in *Escherichia coli* BL21 DE3 cells (Stratagene) with the help of the rare codon plasmid pRARE. An exception presents a *M. jannaschii* Connectase variant with a removable N-terminal $His_6$-tag, which was used for crystallography and cloned into a pET28b vector, instead. Through choice of appropriate PCR primers, the following variants were produced: *M. Mazei* Connectase (MM_2909) was cloned together with a C-terminal $His_6$- (GSHHHHHH), Strep- (GSWSHPQFEK), Myc- (GSEQKLISEEDL) or HA-tag (GSYPYDVPDYA) and as active site mutant Connectase$^{T1A}$ with C-terminal $His_6$-tag; *M. jannaschii* Connectase (MJ_0548) was cloned with a C-terminal $His_6$-tag, as active site mutant Connectase$^{S1A}$ with C-terminal $His_6$-tag, without beta-barrel domain (Connectase$^{\Delta\beta B}$, lacking residues 203-293; C-terminal $His_6$-tag), as active site mutant without beta-barrel domain (Connectase$^{\Delta\beta B\ S1A}$, C-terminal $His_6$-tag) and without NTN-domain (Connectase$^{\Delta NTN}$, lacking residues 1-200; C-terminal $His_6$-tag). Truncated MtrA constructs, *M. mazei* MtrA$^{\Delta 219\text{-}240}$ (MM_1543) with N-terminal Strep-tag (MWSHPQFEKGS) as well as *M. jannaschii* MtrA$^{\Delta 225\text{-}245}$ and MtrA$^{\Delta 174\text{-}245}$ (MJ_0851) with N-terminal $His_6$-tag (MHHHHHHGS), were also generated. In a similar way, based on the gene for mature *C. subterraneum* ubiquitin (Csub_C1474, synthesized by Eurofins), the following N-terminally $His_6$-tagged primary ligation substrates or C-terminally $His_6$-tagged secondary ligation substrates were produced: Ubiquitin fused C-terminally to the *M. mazei* MtrA residues 148-154 (Ub-(5)KD), 148-167 (Ub-(5)KDPGA(10)) or 148-172 (Ub-(5)KDPGA(15)) or N-terminally to a start-methionine plus *M. mazei* MtrA residues 155-167 (PGA(10)-Ub), 155-172 (PGA(15)-Ub) or 155-177 (PGA(20)-Ub). Furthermore, the following genes were synthesized by Biocat: Mature *C. subterraneum* ubiquitin with N-terminal $His_6$-tag and C-terminal *M. mazei* MtrA 148-172 sequence plus Strep-tag ($His_6$-Ub-(5)KDPGA(15)-Strep), with N-terminal *M. mazei* MtrA 155-172 sequence and without affinity tag (PGA(15)-Ub), with N-terminal $His_6$-tag and C-terminal *M. jannaschii* MtrA sequence 154-173 (Ub-(5)KDPGA(10)), with C-terminal $His_6$-tag and N-terminal *M. jannaschii* MtrA sequence 161-178 (PGA(15)-Ub), with a C-terminal GGSLPETGGG modification and $His_6$-tag (Ub-GGSLPETGGG) or with an N-terminal $His_6$-tag following a TEV site for N-terminal glycine exposure (GGG-Ub). Camelid α-ricin single-domain antibody[7] (sdAb), Cyclophilin (CyP) or Glutathione-S-Transferase (GST) sequences were fused to an N-terminal *M. mazei* PGA(10) sequence and a C-terminal $His_6$-tag. Finally, Sortase A (SrtA) and the Sortase A pentamutant (SrtA5*) were synthesized as decribed previously [8], with an N- (SrtA) or C-terminal (SrtA5*) $His_6$-tag and without membrane anchor (i.e. without residues 1-59). All protein sequences and primers used for their generation are listed in the supplementary information.

Cells were grown at 25°C in M9 minimal medium supplemented with 50 µg/ml Se-Met, Leu, Ile, Phe, Thr, Lys and Val for Se-Met labeling, or in lysogeny broth (LB) for all other purposes. Protein expression was induced at an optical density of 0.4 at 600 nm with 500 µM isopropyl-β-D-thiogalactoside. Cells were harvested after 16 h, lysed by French press, and cleared from cell debris by ultracentrifugation. $His_6$-tagged proteins were purified via HisTrap HP columns (20 mM Tris-HCl pH 8.0, 250 mM NaCl, 20 - 250 mM imidazole; all columns obtained from GE Healthcare) and Strep-tagged constructs via HiTrap Streptavidin HP columns (20 mM HEPES-NaOH pH 7.5, 250 mM NaCl, 0 – 2.5 mM desthiobiotin). The $His_6$-tags of the GGG-Ub substrate and the N-terminally tagged Connectase variant for crystallography were then removed using the TEV or Thrombin proteases, respectively. Next, the thermostable *M. jannaschii* and *C. subterraneum* proteins were incubated for 10 min at 80°C and denatured protein removed via centrifugation. Finally, all proteins were applied to a Superdex 75 size-exclusion column (20 mM HEPES-NaOH pH 7.5, 100 mM NaCl, 50 mM KCl, 0.5 mM TCEP). All chromatography steps were performed on an Äkta Purifier FPLC (GE Healthcare) using Unicorn v5.1.0 software. Purified proteins were supplemented with 15% glycerol, flash frozen in liquid nitrogen and stored at -80°C. Connectase and SrtA secondary substrates (PGA(X)-Ub; GGG-Ub) were subsequently analysed via LC-MS (see LC-MS section) to ensure complete removal of the start-methionine.

### *Pulldown and mass spectrometrical analysis*

*M. mazei* Connectase fused to Strep- Myc- or HA-tags was recombinantly expressed in 50 ml LB medium, lysed and cleared from cell debris as described above, except for the use of a different resuspension buffer (buffer A: 20 mM MOPS-NaOH pH 7.1, 150 mM NaCl, 100 mM KCl, 0.15% NP-40). Likewise, 20 g of stationary phase *M. mazei* Go1 cells were resuspended in 50 ml buffer A, lysed and insoluble fractions removed via ultracentrifugation. Magnetic Streptavidin- (87.5 µl), anti-Myc or anti-HA magnetic beads (175 µl each; Thermo) were then incubated for 1 h at room temperature with the above extracts, containing Strep-, Myc- or HA-tagged Connectase, respectively. Afterwards, the beads were washed five times with 1 ml of buffer A and incubated with 15 ml *M. mazei* extract for 1 h at room temperature. After two washes with buffer A, bound proteins were eluted with 40 µl buffer A supplemented with 10 µM desthiobiotin or 2 mg/ml HA- or Myc-peptides, respectively. For mass spectrometrical analysis, bound proteins were separated via SDS-PAGE, following an in-gel tryptic digest (13 ng/µl trypsin, 20 mM ammonium bicarbonate) [9]. LC-MS/MS analysis on a Proxeon Easy nano-LC (Thermo) coupled to an LTQ OrbitrapElite mass spectrometer (Thermo). The data were processed using MaxQuant v1.6.4 [10] and spectra searched against the Uniprot *M. mazei* Go1 proteome (Extended Data, Fig. 3a).

### *Co-elution experiment*

To verify the Connectase-MtrA interaction (Fig. 1a), equimolar (20 µM each in 1 ml) mixtures of either *M. mazei* Connectase-$His_6$ or $Connectase^{T1A}$-$His_6$ and Strep-$MtrA^{\Delta 219\text{-}240}$ were applied on a HisTrap HP 1 ml column (GE Healthcare). After rigorous washing with binding buffer (20 mM Tris-HCl pH 8.0, 250 mM NaCl, 20 mM imidazole), bound proteins were eluted in a single step with increased imidazole concentrations (250 mM) and the obtained fractions analyzed via SDS-PAGE.

### *Liquid chromatography mass spectrometry (LCMS)*

For analysis of the MtrA-Connectase conjugate (Fig. 1b-c), 0.5 g/l $His_6$-tagged *M. mazei* Connectase was incubated with 0.5 g/l Strep-tagged *M. mazei* $MtrA^{\Delta 219\text{-}240}$ in size-exclusion buffer for 20 min on ice. Similarly, for analysis of the Connectase-mediated protein-protein ligation (Extended Data, Fig. 6), 0.5 g/l $His_6$-tagged *M. mazei* Connectase was incubated with 0.5 g/l $His_6$-tagged Ub-(5)KDPGA(10) and 0.5 g/l $His_6$-tagged PGA(10)-Ub. Desalted samples were subjected to a Phenomenex Aeris Widepore 3.6 µm C4 200 Å (100 x 2.1 mm) column, eluted with a 30-80% $H_2O$/acetonitrile gradient over 15 min in the presence of 0.05% trifluoroacetic acid and analyzed with a Bruker Daltonik microTOF. Data processing was performed with Bruker Compass DataAnalysis 4.2 and the m/z deconvoluted with the MaxEnt module to obtain the protein mass.

### *Mass spectrometrical analysis of dimethyl labeled MtrA-Connectase*

To determine free amino groups in *M. mazei* Strep-$MtrA^{\Delta 219\text{-}240}$, Connectase-$His_6$ and their reaction product $MtrA^N$-Connectase, the corresponding bands were excised from an SDS-gel similar to the one shown in Fig. 1a. Following an in-gel tryptic digest (13 ng/µl trypsin, 20 mM ammonium bicarbonate) [9], extracted protein fragments were desalted with C18 StageTips [11] and dimethylated (0.16 % $CH_2O$, 22 mM $NaBH_3CN$, 100 mM TEAB; [12]) with an incorporation rate of 89 - 92%. Due to the use of different isotopes, this procedure resulted in a 28 Da ("light", $MtrA^N$-Connectase), 32 Da ("medium", MtrA) or 36 Da ("heavy", Connectase) mass shifts per dimethylated amino group, respectively. LC-MS/MS analysis was performed subsequently on a Proxeon Easy nano-LC (Thermo) coupled to an LTQ OrbitrapElite mass spectrometer (Thermo). The data were processed using MaxQuant v.1.6.4 [10] and spectra searched against a custom peptide database and the Uniprot M. mazei Go1 proteome (Fig. 1d; Extended Data, Fig. 3b).

### *Light scattering*

Static light-scattering experiments (SEC-MALS; Fig. 2a; Extended Data, Fig. 5a-d) were performed with 50 µl 200 µM *M. jannaschii* $Connectase^{S1A}$-$His_6$, $Connectase^{\Delta\beta B\ S1A}$-$His_6$, $Connectase^{\Delta NTN}$-$His_6$, $His_6$-$MtrA^{\Delta 225\text{-}245}$, $His_6$-$MtrA^{\Delta 174\text{-}245}$ or 1:1 molar mixtures of the respective Connectase-MtrA pairs, using a Superdex S200 10/300 GL gel size-exclusion column (20 mM HEPES-NaOH pH 7.5, 50 mM NaCl, 100 mM KCl) coupled to a miniDAWN Tristar Laser photometer (Wyatt) and a RI-2031 differential refractometer (JASCO). Data analysis was carried out with ASTRA v7.3.0.18 software (Wyatt). The reported values correspond to the mean (± standard deviation) of three independent experiments.

### *Microscale thermophoresis*

For microscale thermophoresis (MST) experiments (Fig. 2b; Extended Data, Fig. 5e-l), peptides based on the *M. jannaschii* MtrA sequence were synthesized by GenScript, with the fluorophore fluorescein isothiocyante (FITC) either N-terminally linked via aminohexanoic acid ((15)KDPGA(5), (15)KDPGA(0)) or an an extra C-terminal lysine ((15)KDPGA(10), (10)KDPGA(10), (5)KDPGA(10), (0)KDPGA(10); see supplementary information). Purified *M. jannaschii* $His_6$-$MtrA^{\Delta 225\text{-}245}$ and $His_6$-

MtrA$^{\Delta 174\text{-}245}$ proteins were fluorescently labeled using the NT-647-NHS labeling kit (Nanotemper). Next, a serial 1:1 dilution of *M. jannaschii* Connectase$^{S1A}$ ranging from nano- to micromolar concentrations was prepared and mixed with 10 nM labeled peptide or 50 nM labeled protein (20 mM HEPES-NaOH pH 7.5, 150 mM NaCl, 50 mM KCl, 0.5 mM TCEP, 0.05% NP40, 0.1 g/l BSA). MST measurements were performed with a Monolith NT.115 (Nanotemper), using various MST power and laser intensity settings to test the general validity of the obtained data. The reported values were obtained at a temperature of 25°C and correspond to the mean (± standard deviation) of three independent experiments. The shown binding curve was fitted to the data, using the NT Analysis 1.5.41 software (Nanotemper).

***Crystal structure determination***

For initial crystallization experiments, *M. jannaschii* Connectase was concentrated to 15 mg/ml in 150 mM NaCl, 0.5 mM TCEP and 20 mM Hepes pH 7.5. As a remnant of the Thrombin cleavage site (see Cloning, Expression and Purification), the employed protein variant had a four-residue modification, GSHM, on the N-terminus. Screening of conditions was performed at 21 °C in 96-well sitting-drop vapor-diffusion plates with 600 nl drops containing equal volumes of protein solution and commercial screening solutions, equilibrated against a 50 µl reservoir. After optimization of conditions, best diffracting crystals were obtained with a crystallization buffer containing 70% MPD and 100 mM Tris pH 8.5. Crystals were loop-mounted directly from the plates and flash-cooled in liquid nitrogen. Diffraction experiments were performed at 100K and a wavelength of 1Å at beamline X10SA in 2009, using a MarCCD 225mm CCD detector. Data were indexed, integrated and scaled using XDS [13], yielding a dataset in space group C2 with a resolution cutoff at 2.3Å (Extended Data, Fig. 5m). Unit cell dimensions and space group led us to expect 2-7 protein molecules in the asymmetric unit (ASU); attempts to solve the structure via molecular replacement using different truncated structures of proteasome beta subunits remained unsuccessful.

We thus reproduced the crystals using a selenomethionine labeled version of the same *M. jannaschii* Connectase protein for anomalous dispersion experiments, which required further adjustment and optimization of conditions to obtain crystals of sufficient quality; best diffracting crystals were obtained at 30 °C with a protein concentration of 6 mg/ml and a crystallization buffer containing 70% MPD and 100 mM HEPES pH 7.5, which were again loop-mounted directly from the plates and flash-cooled in liquid nitrogen. Due to the low lattice symmetry (C2) and the limited possibilities to record highly redundant data on the CCD detector, initial SAD experiments at the Se K-edge yielded only faint anomalous difference signal. This led us to perform 4-wavelength MAD experiments, which yielded data to about 2.8 Å (Extended Data, Fig. 5m), with detectable anomalous differences extending to about 3.8 Å. After indexing, integration and scaling using XDS, we successfully employed SHELXD [14] for heavy atom location, followed by substructure refinement using SHARP [15]. After density modification with Solomon [16], the ARP/WARP secondary structure recognition pipeline [17] could trace the backbone of the four protein molecules in the asymmetric unit to large extents, such that most of the structure could be built subsequently by Buccaneer [18]. At this point, we switched to refine against the higher-resolution native data and completed the structure by cyclic manual modeling with Coot [19] and refinement with REFMAC5 [20] with local NCS restraints (Extended Data, Fig. 5m). The Ramachandran statistics (most favored/additionally allowed/generously allowed) are 94.6%/4.7%/0.4% as assessed with PROCHECK [21].

To obtain a complex structure, an equimolar ratio of Connectase$^{S1A}$ and FITC-Ahx-(15)KDPGA(10) peptide were concentrated to 10.5 mg/ml in 50 mM NaCl, 0.5 mM TCEP and 20 mM HEPES pH 7.5. Crystallization screens were performed as described above. Best diffracting crystals were identified with condition #21 of the NeXtal ProComplex suite (100 mM Na-cacodylate, 15% PEG 4000) as crystallization buffer; prior to loop-mounting and flash-cooling in liquid nitrogen, crystals were briefly transferred to a droplet of crystallization buffer supplemented with 20% glycerol for cryoprotection. Diffraction experiments were performed at 100K and a wavelength of 1Å at beamline X10SA in 2019, using a Pilatus 6M-F hybrid pixel photon counting detector. Data were indexed, integrated and scaled using XDS, yielding a dataset in space group $P2_12_12_1$ with a resolution cutoff at 3.05Å (Extended Data, Fig. 5m). Molecular replacement with MOLREP [22], using the above *M. jannaschii* Connectase structure as a search model, located 2 monomers in the ASU. After first rounds of refinement with REFMAC5, electron density for one peptide per protein monomer became apparent, which was built manually. The structure was completed by manual modeling with Coot and refinement with REFMAC5 with strong NCS restraints (Extended Data, Fig. 5m). According to PROCHECK, the Ramachandran statistics (most favored/additionally allowed/generously allowed) are 90.1%/9.2%/0.0%.

### *Activity assays*

Unless indicated otherwise, all reactions with *M. mazei* proteins were carried out in *M. mazei* reaction buffer (50 mM acetate, 50 mM MES, 50 mM HEPES, 150 mM NaCl, 50 mM KCl, 5 mM TCEP, pH 7.0) and all experiments with *M. jannaschii* proteins conducted in *M. jannaschii* reaction buffer (50 mM MES, 200 mM NaCl, 50 mM KCl, 5 mM TCEP, pH 5.8). The reactions were stopped at the indicated time points (see figures) by addition of 2% SDS and analyzed on 12% NuPAGE SDS-polyacrylamide gels (Thermo).

The time course of the Connectase reaction with just one primary substrate (Fig. 3b, lanes 1-4) was performed by incubating 0.9 µM *M. mazei* Connectase-$His_6$ with 3.7 µM $His_6$-Ub-(5)KDPGA(10) at 37°C. For the recombination experiment (Fig. 3b, lanes 5-10), 3.7 µM PGA(10)-sdAb-$His_6$, PGA(10)-CyP-$His_6$ or PGA(10)-GST-$His_6$ were added in addition and the samples incubated for 10 min before SDS-PAGE analysis.

To determine ligation rates with different recognition motifs (Table 1; Extended Data, Fig. 7), peptides based on the *M. mazei* MtrA sequence were synthesized by GenScript (Supplementary Information). The experiments were then conducted by incubating 25 µM of the indicated primary and secondary substrates with 62.5 nM *M. mazei* Connectase-$His_6$ at 50°C, following SDS-PAGE band quantification of the reaction time course. For the ligation with Ub-(5)KDPGA(15) and PGA(10) / PGA(15) peptides, an additional Strep-tag sequence (indicated by an asterisk) was added C-terminally of the Ub-(5)KDPGA(15) substrate, so that the ligation results in a size shift. After adjusting loaded protein quantities and the Coomassie staining procedure to ensure a linear, concentration-dependent signal increase, three independent experiments were conducted for each substrate combination. Protein bands were quantified with ImageJ 1.52a, assuming that all ubiquitin molecules bind the Coomassie dye in a similar manner. The obtained data showed that the observed product formation in reactions with equimolar educts correlated with the quotient between the current and maximum product formation:

$$Observed\ Ligation\ \% = \int_{t\_start}^{t\_end} [Maximum\ rate] * (1 - \frac{Product\ concentration\ at\ t}{Maximum\ product\ concentration}) * \mathrm{dt}$$

where the observed ligation % is the molar quotient between products and educts. This relationship was found to describe the empirical data best (see fits in Extended Data, Fig. 7-9) and used to approximate the maximum rate parameters given in Fig. 4b (mean of three experiments ± standard deviation).

Kinetic parameters (Fig. 4c, Extended Data, Fig. 8) of a Connectase reaction were determined by incubating the indicated concentrations of $His_6$-Ub-(5)KDPGA(15) and PGA(15)-Ub-$His_6$ at 50°C with 400x lower concentrations of *M. mazei* Connectase-$His_6$. The maximum rate parameters of each experiment were determined as described above and used in a Michaelis-Menten plot. The kinetic parameters were determined based on a data fit using the Michaelis-Menten model.

The role of the C-terminal beta-barrel domain (Extended Data, Fig. 9) was studied by recording and analyzing time courses of *M. jannaschii* Connectase or Connectase$^{\Delta\beta B}$ ligations as described above, except for the use of *M. jannaschii* MtrA-derived $His_6$-Ub-(5)KDPGA(10) and PGA(15)-Ub-$His_6$ substrates and a reaction temperature of 85°C.

For a comparison with Sortase A (SrtA) and the Sortase A pentamutant (SrtA5*; Fig. 4a; Extended Data, Fig. 10), the data gathered in the kinetic analysis (see above) was compared with analogous SrtA5*/SrtA ligations using low (3.1 µM) or high (100 µM) substrate concentrations. The SrtA5*/SrtA reactions were conducted in the same manner as the Connectase reaction, instead for the use of Ub-GGSLPETGGG-$His_6$ and GGG-Ub substrates, a reaction temperature of 37°C, a different buffer system (50 mM Tris-HCl pH 7.5, 150 mM NaCl, 10 mM CaCl2; [23]) and higher enzyme concentrations as indicated. A data fit for SrtA5*/SrtA reactions was not possible due to the competing catalytic activities (Hydrolysis, Ligation).

A potential Connectase hydrolase activity (Fig. 4d) was investigated by incubating a mix of 25 µM $His_6$-Ub-(5)KDPGA(15)-Strep and 25 µM PGA(10) with either 25 nM or 25 µM *M. mazei* Connectase-$His_6$ at 50°C, following SDS-PAGE analysis.

For the ligation within crude extracts (Fig. 4e), Strep-Ub-(5)KDPGA(10)-$His_6$ and PGA(15)-Ub without affinity tag were recombinantly expressed in *E. coli*. The lysate of a 2 l culture was prepared as described above, except for using a different resuspension buffer (50 mM acetic acid, 50 mM MES, 50 mM HEPES, 100 mM NaCl, 50 mM KCl, 10 mM Imidazole, 5 mM $MgCl_2$, 50 µg/ml DNAse (Applichem) and c0mplete protease inhibitor (Roche) pH 7.0). For the ligation, equal volumes of each cell lysate were mixed with 0.09 g/l *M. mazei* Connectase-$His_6$, which corresponds to a molar enzyme:substrate ratio of roughly 1:30. After incubation for 15 min at 37°C, the pH was then adjusted to 8.0 and the mixture applied on a HisTrap FF $Ni^{2+}$-NTA column in series with a HiTrap streptavidin column (GE). Following rigorous washing (20 mM Tris, 250 mM NaCl, pH 8), the reaction product was eluted with 2.5 mM desthiobiotin and, in a second step, $His_6$-tagged educts with 2.5 mM desthiobiotin and 250 mM imidazole.
